# Quantitative analysis of morphogenesis and growth dynamics in an obligate intracellular bacterium

**DOI:** 10.1101/2022.12.26.521939

**Authors:** Wanda M. Figueroa-Cuilan, Oihane Irazoki, Marissa Feeley, Erika Smith, Trung Nguyen, Felipe Cava, Erin D. Goley

## Abstract

Obligate intracellular bacteria of the order Rickettsiales include numerous arthropod-borne human pathogens. However, our understanding of the basic biology of *Rickettsia* species is limited by technical challenges imposed by their obligate intracellular lifestyle. To overcome this roadblock, we developed quantitative methods to assess the cell wall composition, intracellular growth, and morphology of *Rickettsia parkeri*, a human pathogen in the Spotted Fever Group of the *Rickettsia* genus. Analysis of the cell wall composition of *R. parkeri* revealed unique features including a high M3 monomer fraction and absence of LD-crosslinks. Using a novel fluorescence microscopy approach, we quantified the cell morphology of *R. parkeri* in live host cells and found that bacterial morphology is maintained stably during exponential growth in two different epithelial cell lines. To assess population growth kinetics in a high-throughput and high-resolution manner, we developed an imaging-based growth assay and applied this to determine the growth rate of up to 24 infected cultures at a time. We also sought to gain insight into the cell cycle regulation of *R. parkeri*. To this end, we developed methods to quantify the fraction of the population preparing to divide as well as those undergoing active constriction. These approaches permitted a quantitative analysis of cell cycle status across a population of *R. parkeri.* Finally, as a proof of concept, we applied the above tools to quantitatively determine how MreB, a bacterial actin homolog, contributes to the growth and morphogenesis of *R. parkeri*. Inhibition of MreB with the small molecule MP265 led to cell rounding and slowed growth, suggesting that MreB is required for the growth and shape maintenance of *R. parkeri*. Collectively, we developed a toolkit of high-throughput, quantitative tools to understand intracellular growth and morphogenesis of *R. parkeri* that is translatable to other obligate intracellular bacteria.

**AUTHOR SUMMARY:** The obligate intracellular lifestyle of members of the bacterial order Rickettsiales, which includes important human pathogens, has hindered our progress in understanding their biology. Here we developed and applied high-throughput, quantitative tools to analyze essential features of rickettsial cell biology such as morphology and growth in living host cells. By applying these tools in a proof of concept, we showed that the bacterial actin homolog, MreB is required for the regulation of rod shape and intracytoplasmic growth.

## INTRODUCTION

The order Rickettsiales includes medically relevant, emerging bacterial pathogens that grow exclusively inside a host cell and may be transmitted to mammals by ticks and other arthropods. Examples of important human rickettsioses are Spotted Fever, Typhus Fever Rickettsioses, and Scrub Typhus. Early clinical manifestations are non-specific between rickettsioses, and include fever, headache, rash, confusion, and myalgia (1). *Rickettsia* species exhibit a broad host range; however, all known human pathogenic rickettsioses are transmitted by arthropods (2–5). Spotted Fever Group (SFG) *Rickettsia* are emerging pathogens that underlie ∼5,000-6,000 diagnosed cases in the United States (USA) annually. The fatality rate reported for the most severe rickettsiosis in the USA, Rocky Mountain Spotted Fever (RMSF) caused by *Rickettsia rickettsii*, is 5-10% with treatment (5) and up to 55% without treatment (6). In fact, the Centers of Disease Control (CDC) considers RMSF the “most deadly tick-borne disease in the world” (7). Thus, a deeper understanding of essential mechanisms that allow pathogenic *Rickettsia* to achieve successful infection in human cells is needed.

SFG Rickettsia include *R. rickettsii* (RMSF)*, R. parkeri* (maculatum disease), and *Rickettsia* species 364D, among others. *R. parkeri* rickettsiosis closely resembles RMSF, but the clinical manifestations are less severe. *R. parkeri* has therefore emerged as a valuable model for the study of pathogenic SFG *Rickettsia* (8–11). Like all *Rickettsia*, *R. parkeri* has undergone reductive evolution, which has resulted in the loss of key metabolic enzymes and a strict dependence on a eukaryotic host for growth (12). Indeed, *Rickettsia* species have been predicted to depend on a eukaryotic host for an estimated 50 essential metabolites (13). In addition, pathways impacted by reductive evolution include those required for the replication of well-studied species like *Escherichia coli* and *Caulobacter crescentus*, such as peptidoglycan (PG) cell wall synthesis, cell division, and cell separation (13, 14). Streamlining of these pathways suggests that *R. parkeri* and other *Rickettsia* species accomplish growth and/or division through alternative or adaptive mechanisms.

Growth and division of most bacteria, including *R. parkeri*, requires synthesis and remodeling of the PG cell wall. The PG consists of glycan strands of repeating N-acetylglucosamine (GlcNAc), N-acetylmuramic acid (MurNAc) sugars cross-linked to each other by short peptides. PG is found in almost all bacteria and plays a fundamental role in the growth and survival of bacterial cells as it maintains envelope integrity against turgor pressure (15). PG is also necessary and sufficient to preserve cell shape (16–20). Conversely, changes in cell shape that occur during growth and division require enzymatic remodeling of PG, which is mediated by a host of tightly regulated PG metabolic enzymes (18,21–23).

In most rod-shaped bacteria, two main PG metabolic complexes mediate growth and division: the elongasome and divisome, respectively (24). The elongasome is a morphogenetic complex that is essential for growth in many rod-shaped organisms and requires MreB, an actin homolog. MreB assembles into short filaments roughly perpendicular to the long axis of the cell that serve as a platform for lateral insertion of cell wall by PG synthases (25, 26). In preparation for division, the divisome complex is directed to mid-cell via the tubulin homolog, FtsZ. Upon polymerization, FtsZ directly and indirectly recruits dozens of essential and accessory proteins to the mid-cell to ultimately direct PG synthesis for inward constriction of the cell envelope and PG hydrolysis during cell separation (27, 28) . Despite decades of intense research in free-living model systems, the mechanisms by which *R. parkeri* and other *Rickettsia* species accomplish growth and division are unclear. This is largely because the obligate intracellular lifestyle of *R. parkeri* has limited the application of genetic and imaging tools to understand these important pathogens (29, 30).

Here, we set out to establish a foundational understanding of *R. parkeri* morphology, growth, and cellular organization as a model for SFG *Rickettsia* cell biology. To do this, we first established rigorous, quantitative methods to assess PG composition, morphology, and growth. We applied these to visualize a morphogenetic complex in *R. parkeri* and to quantitatively analyze the role of the elongasome in morphogenesis and growth. This work establishes a basic understanding of cell wall biochemistry and morphogenesis of an understudied human pathogen, and provides tools to probe mechanisms of rickettsial growth in the intracellular environment that are translatable to other obligate intracellular bacteria.

## RESULTS and DISCUSSION

### PG Composition of *R. parkeri* Represents a Distinct Class

To begin dissecting the pathways that contribute to the intracellular growth and morphogenesis of *R. parkeri* in an unbiased fashion, we performed a muropeptide composition analysis of its PG sacculus (31). This approach allows the identification of cell wall components (muropeptides) and their abundance, which reflect major cell wall enzymatic activities under the conditions tested (32). Of note, isolating enough cell wall from any Rickettsiae grown in host cells is challenging due to the extremely low PG yield (33). To circumvent this challenge, we isolated PG from *R. parkeri* grown in two large-scale biological replicates of infected African green monkey kidney epithelial Vero cells. These were then subjected to enzymatic digestion and analyzed by ultraperformance liquid chromatography (UPLC), and tandem mass spectrometry (MS/MS). In agreement with other *Rickettsia* species muropeptide composition analyses, including that of *R. typhi* (33, 34), we detected muropeptides containing *meso*-diaminopimelic acid (DAP) as the third amino acid, which is typical of Gram-negative PG (19) (**Fig 1**). UPLC-MS data showed that the largest monomeric structure of the *R. parkeri* PG is the disaccharide pentapeptide GlcNAc-(β 1→4)-MurNAc-(L)-Ala-(D)-Glu (γ)-*meso*-DAP-(D)-Ala-(D)-Ala (or M5). Importantly, we were unable to detect muropeptides in uninfected Vero cells by UPLC-MS suggesting that these muropeptides are specific to *R. parkeri* (**Fig 1A**). We conclude that the *R. parkeri* PG contains the canonical monomer for Gram-negative bacteria.

**Fig 1.**
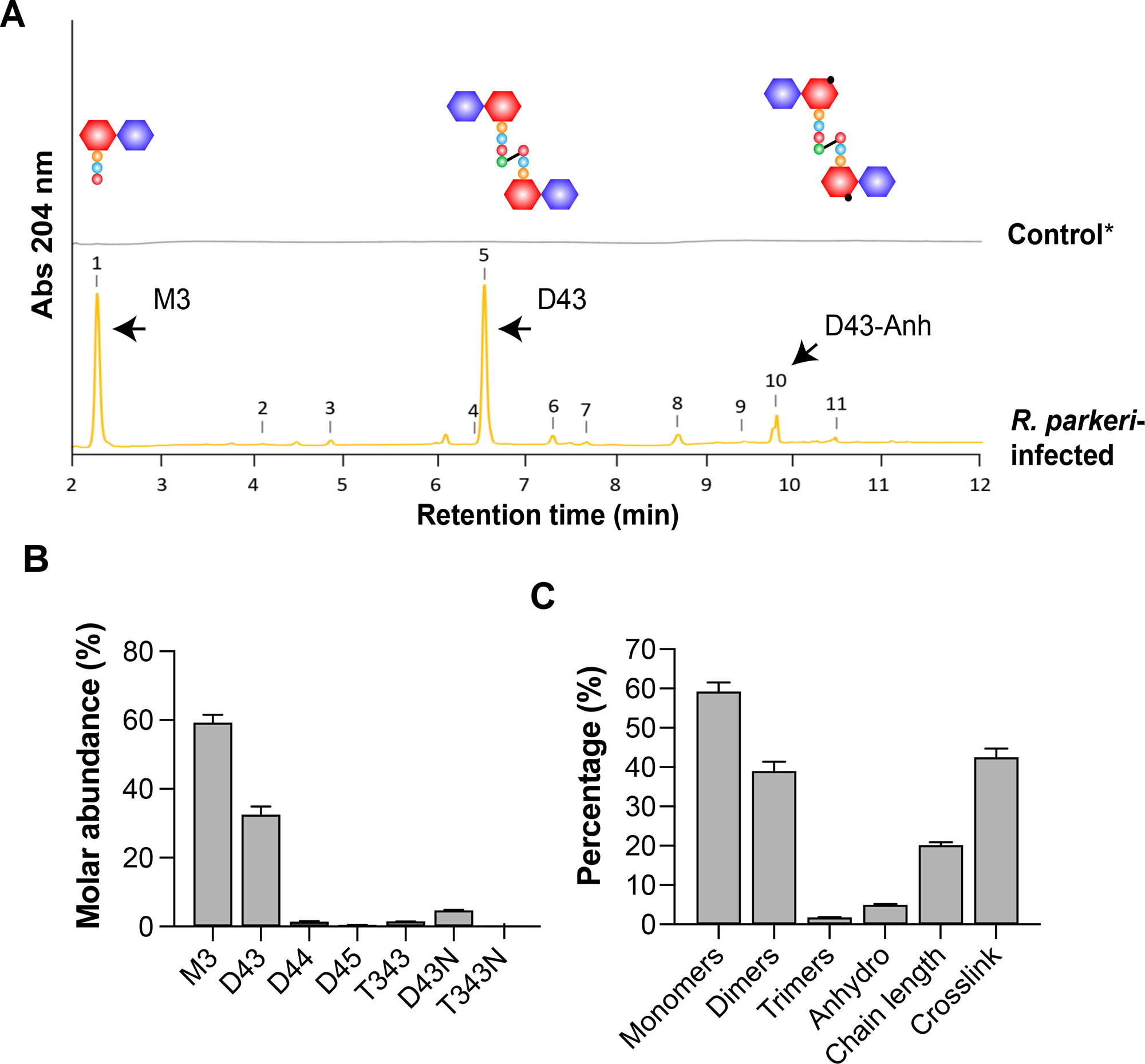
Characterization of *R. parkeri* peptidoglycan by UPLC and mass spectrometry. **(A)** Representative chromatogram of *R. parkeri* PG. The most abundant muropeptides are indicated with arrows and structures are drawn. *Control indicates uninfected Vero cell. **(B-C)** Relative muropeptide molar abundance and the percentage of each muropeptide detected in *R. parkeri* PG.

To gain insight into which PG enzymes may be active during intracellular PG synthesis and remodeling, we characterized the *R. parkeri* PG cell wall chemical composition in detail by UPLC. We identified 11 different muropeptides, the MS and retention times of which matched the expected values for previously described DAP-type PG (**Fig 1A**, **Supplemental Table 1**). Among the identified muropeptides, most of the *R. parkeri* PG subunits appeared as uncrosslinked monomers (∼60% of the relative abundance), specifically disaccharide tripeptides (M3s) (**Fig 1BC**, **Supplemental Table 1**). This suggests that enzymes that cleave stem peptides between the m-DAP and D-Ala positions are highly active during intracellular growth. Consistent with previous bioinformatic analyses that indicated the absence of LD-transpeptidases encoded by *R. parkeri*, our UPLC analysis showed no detectable LD-crosslinked muropeptides (**Fig 1B**), implicating LD-carboxypeptidases after DD-carboxypeptidation and/or LD-endopeptidases in generation of M3s (14).

Of note, we attempted to visualize the patterning of PG insertion in *R. parkeri* by incubating infected cells with Fluorescent D-Amino Acids (FDAAs) (35), but this proved unsuccessful. FDAAs are incorporated into PG in the periplasm, through DD- or LD-transpeptidase activity (36). The lack of LD-transpeptidation in *R. parkeri* may partly explain the lack of FDAA staining in *R. parkeri*. In addition, the high M3 abundance we observed, reflecting LD-carboxypeptidase and/or LD-endopeptidase activity, would result in the removal of FDAAs incorporated by LD- or DD-transpeptidases, which are incorporated into the 4^th^ or 5^th^ position of the peptide stem, respectively (36).

The genome of *R. parkeri* encodes a putative periplasmic LD-carboxypeptidase homolog, LdcA (MC1_RS02825), which we hypothesize is of primary importance in the cleavage of peptide stems to generate tripeptide-containing products. The biological significance for the high M3 abundance in *R. parkeri* remains to be uncovered; however, it has been previously shown that the DAP-type PG cell wall or DAP-containing PG fragments are sensed by human intracellular PG receptor NOD1 (hNOD1) (37–39). In *N. gonorrhoeae,* the periplasmic LD-carboxypeptidase LdcA releases tripeptide stems which are sensed by human intracellular PG receptor NOD1 (hNOD1) (37). It remains to be determined whether the abundant tripeptides of *R. parkeri* PG are released to the host cytosol and if so, if these fragments are involved in its pathogenesis.

The second and third most abundant muropeptides identified in our analysis were the dimers D43 (∼32% relative abundance) and D43 anhydromuropeptide (∼5% relative abundance), which are indicative of DD-transpeptidase and lytic transglycosylase (LT) activity, respectively (**Fig 1AB**, **Supplemental Table 1**). *R. parkeri* encodes homologs for the bifunctional penicillin binding protein (PBP) PBP1a (MC1_RS06310), the elongation-specific TPase PBP2 (MC1_RS04355), and the division-specific TPase FtsI (MC1_RS04370). The specific contributions of PBPs to *R. parkeri* intracytoplasmic growth remain to be determined, but these factors are likely responsible for formation of DD-crosslinks. We also identified putative homologs of the LTs RlpA (MC1_RS02765) and MltG (MC1_RS02260) and two additional ORFs annotated as LTs (MC1_RS04005 and MC1_RS02815). The roles of these LTs in the *R. parkeri* growth, division, and cell separation remain to be determined, but we predict that they contribute to formation of the anhydromuropeptides observed. The abundance of anhydromuropeptides can be used to calculate the average length PG strands, which was found to be ∼20 disaccharide units for *R. parkeri*. This is similar to well-studied Gram-negative organisms such as *E. coli,* in which the average glycan chain length is 17.8-25.8 disaccharide units (40).

Based on the principal features of the *R. parkeri* PG (high M3 monomer, high D43 dimer, no LD-crosslinking), and compared to other alphaproteobacterial species, we conclude that *R. parkeri* PG represents a distinct class from those previously described (41). The unique features of *R. parkeri* PG may hint at specific evolutionary adaptations shaping the regulation of *R. parkeri* growth and, possibly, pathogenesis. These data establish a baseline for future work aimed at understanding the role of particular enzymes or environmental pressures shaping PG chemistry, as well as the role of specific PG features in the pathogenesis of *R. parkeri*.

### Quantitative evaluation of morphology of *R. parkeri* in host cells

Muropeptide composition analysis provided fundamental insights into PG synthesis and remodeling of *R. parkeri*. We next sought to develop tools that would allow us to quantitatively probe cell shape of live *R. parkeri* cells to permit the dissection of the pathways and parameters that influence morphogenesis. Because of the obligate intracellular nature of *R. parkeri,* and its short reported extracellular viability window (∼30 min), we aimed to perform *in vivo* analysis of morphology in live human lung epithelial cell lines (A549). To do this, we leveraged a *R. parkeri* strain producing cytoplasmic green fluorescent protein (Rp-GFPuv) and reasoned that the shape of the cytoplasm reflects the shape of the cell. Using this approach, we could therefore determine cell morphology by imaging cytoplasmic GFPuv (**Fig 2A**). We found that the GFPuv fluorescence revealed cell shape with good resolution (**Fig 2B**).

**Fig 2.**
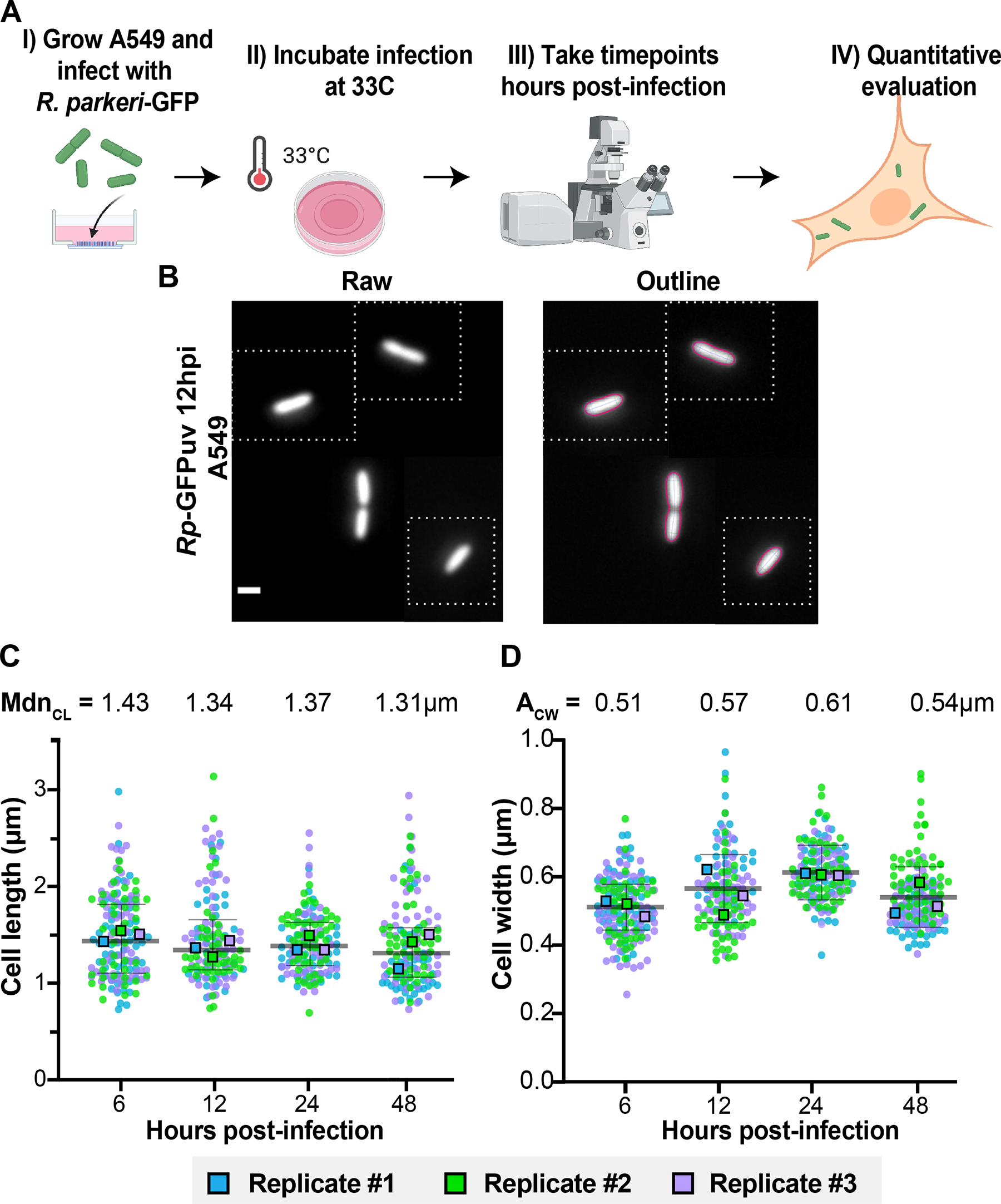
Quantitative evaluation of the morphology of *R. parkeri* in human lung epithelial (A549) cells. **(A)** Strategy for the quantitative evaluation of the morphology of *R. parkeri* producing GFPuv (Rp-GFPuv) in A549 cells. Briefly, human lung epithelial cells (A549) were grown in 35 mm MatTek dishes (Fig 2A, step I), infected with Rp-GFPuv (Fig 2A, step II), and imaged at 6, 12, 24, 48 hours post-infection (Fig 2A, step III). Images were analyzed using MicrobeJ (Fig 2A, step IV). **(B)** Composite of raw Rp-GFPuv images obtained via epifluorescence microscopy (12 hpi) used to detect cell outlines (“Outline”) using MicrobeJ software. Representative cells (bounded by white boxes) were selected from different images to make a composite. Scale bar = 1 µm. **(C)** Cell length distributions of live Rp-GFPuv imaged at indicated hours post infection (hpi) and the middle line represents the global median cell length (Mdn_CL_). **(D)** Cell width distributions of live Rp-GFPuv imaged at indicated hours post infection (hpi) and the middle line represents the global average cell width (A_CW_). Colored dots indicate single cell measurements from three independent biological replicates, whereas the colored boxes represent independent biological replicate Mdn_CL_ or A_CW_. Error bars represent the interquartile range (IQR). N ∼150 cells per timepoint. A Kruskal Wallis with Dunn’s post-test was performed to compare the cell length for all the time points compared to each other, whereas a Welch and Brown-Forsythe ANOVA test with a Dunnett’s post-test was used to compare the cell widths for all the time points to each other. No statistical significance was found comparing cell length or cell width across time points.

The *R. parkeri* intracellular life cycle begins with internalization into host cells and rapid escape from the phagocytic vacuole. Unlike other obligate intracellular bacteria such as the *Chlamydiales,* which grow and divide inside membrane-bound vacuoles or inclusions (42), *R. parkeri* grow and divide freely in the cytoplasm of host cells, and this is followed by actin-mediated cell-to-cell spread (10,11,30). We therefore imaged Rp-GFPuv at 6, 12, 24, and 48 hours post infection (hpi) in A549 cells to reflect pre-, early, and mid-exponential intracytoplasmic growth stages. We used the software MicrobeJ to define the outline of the cells from the GFPuv fluorescence signal (**Fig 2AB**) (11). Using this approach, we measured cell length and width of hundreds of Rp-GFPuv cells at each time point and constructed Superplots (43), where independent biological replicates are represented in different colors (**Fig 2C**). Our analysis revealed a pre-exponential (∼6 hpi) median cell length (Mdn_CL_) of 1.43 µm. As the cells entered early- to mid-exponential phase, the Mdn_CL_ slightly decreased (Mdn_CL_ at 12, 24, and 48hpi = 1.34, 1.37, and 1.31 µm, respectively), but this difference was not statistically significant (**Fig 2C**). Using the same strategy, we calculated the Rp-GFPuv average cell width (A_CW_), which was maintained stably over the time course of infection (A_CW_ after 6, 12, 24, and 48hpi = 0.51, 0.57, 0.61, and 0.54 µm, respectively) (**Fig 2D**).

We next asked whether the morphology of *R. parkeri* varies in different host epithelial cells. To do this, we infected Vero cells with Rp-GFPuv and applied our epifluorescence microscopy strategy to image and quantify live intracellular bacteria at 6, 12, 24, and 48 hpi (**Supplemental Fig 1**). We found that the Mdn_CL_ in Vero cells was ∼1.3 µm throughout infection (Mdn_CL_ 6, 12, 24 and 48hpi = 1.32, 1.25, 1.35, 1.34 µm, respectively) and the A_CW_ in Vero cells was ∼0.5 µm (A_CW_ 6, 12, 24 and 48hpi = 0.48, 0.47, 0.53, and 0.54 µm, respectively) (**Supplemental Fig 1AB**). Previous electron microscopy morphological analysis of *R. parkeri* grown in Vero cells showed a maximum cell length of ∼1.5 µm and an maximum cell width of ∼0.5 µm (44), in good agreement with our high-throughput, fluorescence-based approach.

Because epifluorescence microscopy measurements may be affected by the three-dimensional diffraction pattern of light emitted, we validated our fluorescence strategy by comparing it to quantification of morphology of fixed *R. parkeri* imaged by phase contrast microscopy, a commonly used method in the field of bacterial cell biology (**Supplemental Fig 2A**). To do this, Vero cells were infected with *R. parkeri* or Rp-GFPuv for 48 h, quickly fixed and imaged extracellularly on agarose pads, and quantitatively evaluated in MicrobeJ (**Supplemental Fig 2AB**). We obtained similar values for median cell length (Mdn_CL_ 48hpi = ∼1.4 µm) and width (A_CW_ 48hpi = 0.55 µm) using phase contrast imaging of *R. parkeri* (**Supplemental Fig 2CD**) as with epifluorescence (**Supplemental Fig 1**). Together, our fluorescence-based morphology analysis provides fundamental information about the shape parameters of this enigmatic group of bacteria and provides a robust, high-throughput method for quantification of morphology of intracellular live or fixed *R. parkeri*. Using this method, we conclude that *R. parkeri* cell shape is maintained stably in A549 and Vero cells throughout the early and mid-exponential phases of growth. The ability to reliably quantify morphology of hundreds or thousands of individual bacteria will enable studies into, for example, the reported pleiomorphism of a number of *Rickettsia* species in different growth conditions (45).

### Cell division in *R. parkeri*

Having established a method for quantification of the morphology of *R. parkeri*, we next sought to explore the mechanisms and regulation of *R. parkeri* morphogenesis. As a first step, we focused on cell division, which is mediated by the divisome. We noted from our imaging of Rp-GFPuv the appearance of actively dividing cells, which were identified as those cells with a decrease in GFPuv signal near mid-cell (**Fig 3A**). Demograph analysis of GFPuv fluorescence in cells at 24 hpi revealed that actively dividing cells comprise the longest ∼10% of cells in the population, as expected (**Fig 3B, boxed area**). To understand the impact of infection stage on *R. parkeri* cell division, we quantified the percent of constricting cells in the population at different times post-infection (6, 12, 24, and 48 hpi). We found that the percentage of the population actively undergoing cell division decreased by ∼50% as the cells entered mid-exponential phase and became less variable across replicates (% constriction at 6 hpi = 16.6% in comparison to 48 hpi = 8.35%, respectively) (**Fig 3C**). These data raise the possibility of infection-stage specific input into cell cycle progression of *R. parkeri*.

**Fig 3.**
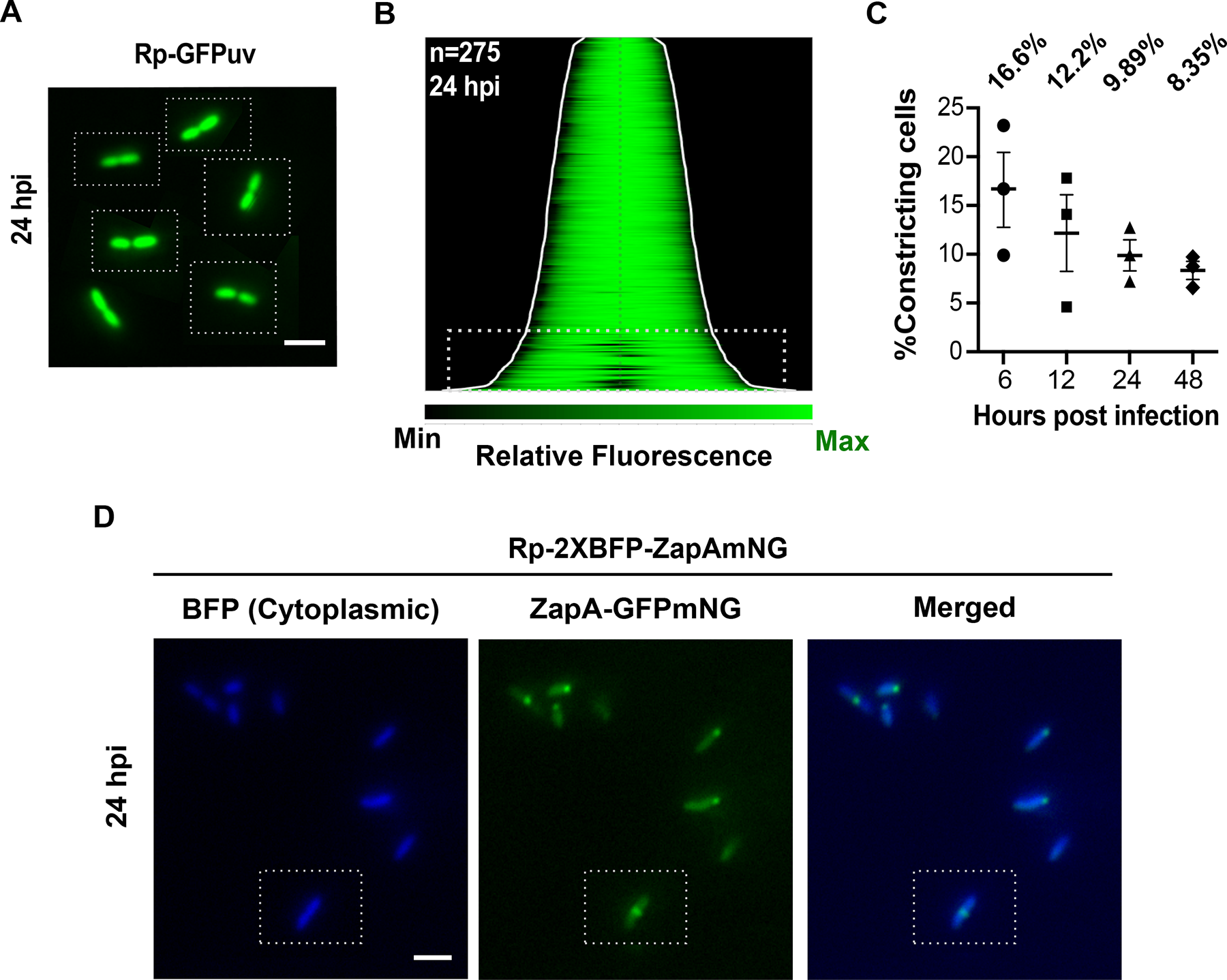
Imaged-based evaluation of *R. parkeri* cell division. **(A)** Composite image of actively dividing intracellular Rp-GFPuv in A549. Representative cells (bounded by white boxes) were selected from different images to make a composite. Scale bar = 2 µm **(B)** A demograph of *R. parkeri* cells producing cytoplasmic GFPuv revealed pre-divisional cells (boxed area). **(C)** Percentage of constricting cells in the population at 6, 12, 24, and 48 hours post infection. **(D)** Spatiotemporal localization of ZapA-mNG on live *R. parkeri* cells expressing cytoplasmic BFP at 24 hpi. Scale bar = 2 µm.

We next sought to visualize the division machinery, itself, in *R. parkeri.* To do this, we focused on imaging a fluorescent fusion to ZapA. ZapA binds to the master regulator of division, FtsZ, at the earliest stages of assembly of the divisome and reliably reports on the localization of the divisome in other bacteria (46–48). We created a *R. parkeri* strain bearing a plasmid encoding *zapA-mNeonGreen* (mNG) driven by the P*_zapA_* promoter (**Fig 3D**). Two copies of the gene encoding a cytoplasmic blue fluorescent protein (TagBFP) driven by P*_ompA_* were included on the same plasmid to mark cells. We found that *R. parkeri* ZapA-mNG displayed polar or mid-cell localization in live cells (**Fig 3D**). This localization pattern is consistent with what has been observed in other alphaproteobacteria, such as *Caulobacter crescentus* (47). To our knowledge, this is the first example demonstrating the subcellular localization of a fluorescent fusion protein in any live rickettsial species. By applying the above tools, we were able to track the cell cycle status of *R. parkeri* across a population of live intracellular bacteria, which will enable future studies aimed at determining how genetic, environmental, and host factors influence the cell cycle and growth of *R. parkeri*.

### Development of a fluorescence-based population growth assay

In addition to quantifying morphology, we sought to develop a high-throughput, high-temporal-resolution growth assay for *R. parkeri*. Measuring population growth kinetics of rickettsial species typically requires quantifying plaque-forming units (PFUs) or genome equivalents over the course of an infection, each of which are labor-intensive, low-throughput methods. To overcome those limitations, we developed a fluorescence-based imaging growth assay in a temperature- and CO_2_-controlled imaging plate reader. We infected A549 cells with Rp-GFPuv for 24 h and imaged GFP fluorescence every 3 h over ∼55 h in a plate reader (**Fig 4A**). We observed an increase in the Rp-GFPuv intensity over time (**Fig 4B**, **Supplemental Videos 1,2**). Next, we plotted the total GFP intensity and used these data to calculate the doubling times for each curve (t_D_). We found that the t_D_ of Rp-GFPuv in A549 cells was ∼4:32h ± 0:28 h (**Fig 4C**).

**Fig 4.**
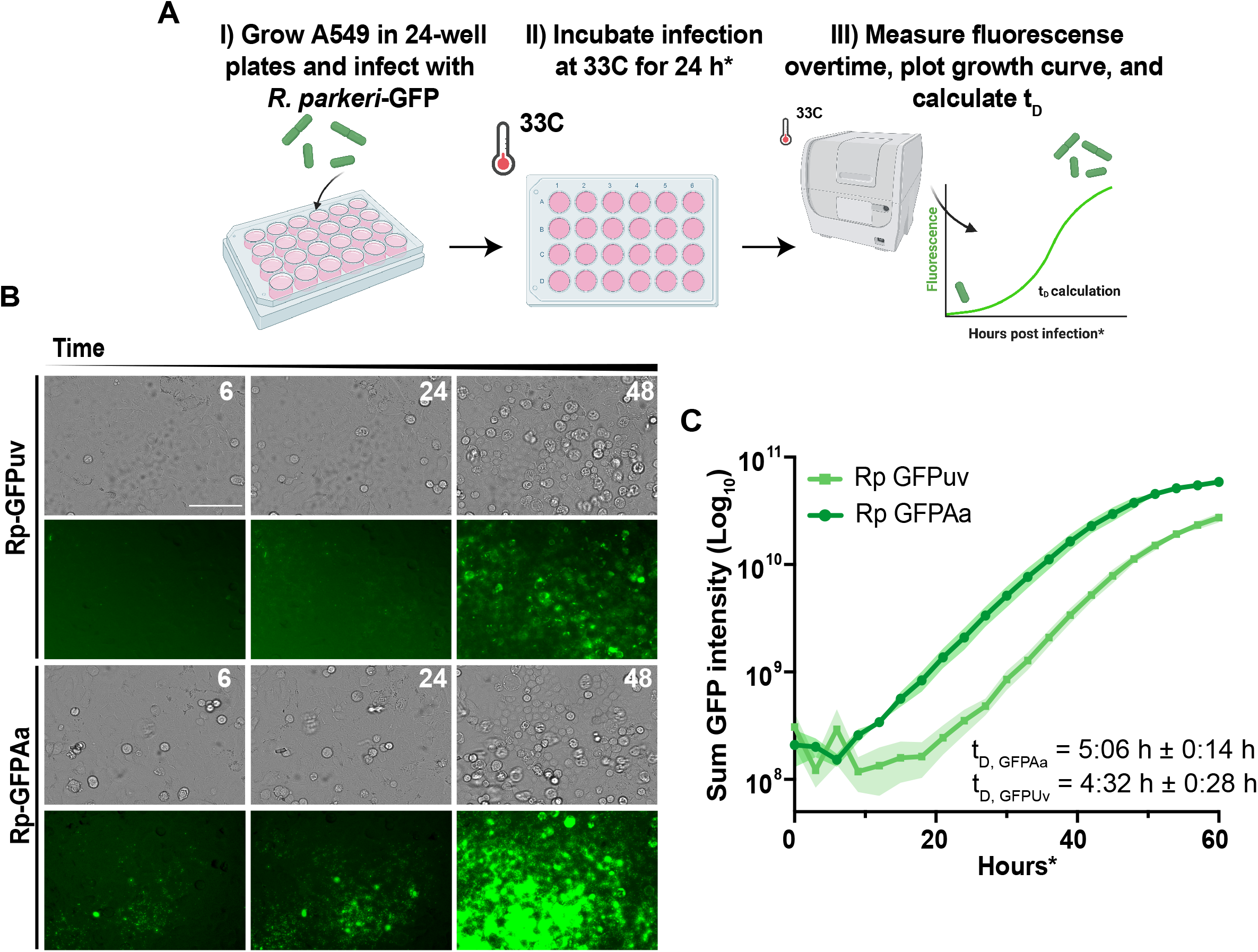
Fluorescence-based quantification of intracellular growth dynamics of *R. parkeri* in A549 cells. **(A)** Strategy for the quantitative evaluation of the intracellular growth of *R. parkeri* in A549 cells. **(B)** Phase-contrast and GFP images taken with the BioTek Cytation1 at 24, 48, 54 hours post-imaging. Scale bar = 100 µm **(C)** Semi-log plot of GFP intensity over time for A549 cells infected with Rp-GFPuv or Rp-GFPAa. t_D_ indicates the doubling time of Rp-GFPuv or Rp-GFPAa.

We noted that it took some time to detect GFPuv above background in our assay. To increase the sensitivity of this assay, we generated a *R. parkeri* strain producing cytoplasmic AausFP1 (Rp-GFPAa), the brightest GFP reported to date (49). GFPAa has a high quantum yield and can be excited with low light, enabling increased sensitivity and long time-lapse fluorescence imaging with limited phototoxicity. Using the same strategy as for Rp-GFPuv (**Fig 4A**), we infected A549s with Rp-GFPAa for 24 h and imaged GFP fluorescence for ∼55 h. We found that Rp-GFPAa fluorescence was significantly brighter, allowing us to detect growth earlier, than Rp-GFPuv (**Fig 4B**, **Supplemental Videos 3,4**) and calculated a t_D_ of 5:06 h ± 0:14 h (**Fig 4C**). To expand our analysis, we also determined the growth dynamics of *R. parkeri* in Vero cells (**Supplemental Fig 3**). We found that the Rp-GFPAa and Rp-GFPuv grow with similar kinetics in Vero cells (t_D_ ∼ 5.5 h for both) (**Supplemental Fig 3B**). We note that we tested our approach monitoring growth of 24 independent infections in the same experiment, enabling high-throughput growth analysis in future studies.

To validate our fluorescence-based growth curve approach, we quantified growth using plaque assays by determining plaque-forming units (PFU)/mL over time to calculate t_D_ for both Rp-GFP strains in Vero host cells. Importantly, our results showed similar t_D_ using PFU growth curves (∼5-6 h) as in our fluorescence-based growth assay (**Supplemental Fig 3C**).

Finally, to assess the impact of expressing the ZapA-mNG fusion protein on the growth kinetics of *R. parkeri*, we applied our fluorescence-based approach to monitor the growth of *R. parkeri* cells producing both cytoplasmic TagBFP and ZapA-mNG from the same plasmid in comparison with cells producing only cytoplasmic TagBFP (**Supplemental Fig 4, Supplemental Videos 5-8**). After imaging BFP fluorescence every 3 h over ∼55 h, we found that the growth kinetics of cells expressing ZapA-mNG (t_D_ of 5:00 h ± 0:48 h) were similar to growth of cells only expressing cytoplasmic TagBFP (5:55 h ± 0:27 h).

Together, these data demonstrate the utility of our fluorescence-based growth assay in determining the growth dynamics of intracellular bacteria such as *R. parkeri* in a high-throughput manner and at high temporal resolution.

### Inhibition of MreB causes cell rounding and slow growth

With assays for quantitative evaluation of growth and morphology in hand, we sought to apply these tools to assess the contribution of molecular machinery predicted to impact morphogenesis in *R. parkeri.* In many rod-shaped bacteria, the elongasome directs insertion of PG along the cylinder of the cells, resulting in cell elongation during growth and maintaining constant cell width. MreB is an actin homolog that acts as a scaffold for PG synthesis enzymes of the elongaosome (26,50–52). Using BLAST, we found that *R. parkeri* encodes putative homologs of all components of the elongasome found in well-characterized Gram-negative species (MreB (MC1_RS06060), MreC (MC1_RS06055), MreD (MC1_RS04360), RodZ (MC1_RS05965), PBP2 (MC1_RS04355), and RodA (MC1_RS01930)) (24, 53). This suggests that the elongasome functions to maintain *R. parkeri* rod shape. To test this hypothesis, we used a pharmacological inhibitor of MreB (MP265) coupled with our morphological and growth assays (54). We infected A549 cells with Rp-GFPuv, treated the infections with 25, 50, or 100 µM MP265, and performed fixed epifluorescence microscopy and MicrobeJ analysis of cell shape (**Fig 5**). We found that treatment with MP265 led to a significant increase in cell width in a dose-dependent manner (A_CW_ 25, 50, and 100 µM = 0.53, 0.70, and 0.69 µm) (**Fig 5ABC**). Moreover, principal component analysis of cell shape using Celltool (55) enabled a more comprehensive analysis of how shape varied between control and MP265-treated cells (**Supplemental Fig. 5**). We found that when Shape Modes 1 (roughly reflecting cell length) and 2 (roughly reflecting cell width) were plotted against each other, Rp-GFPuv treated with 50 or 100 µM MP265 segregated to the bottom two quadrants of the plot, reflecting their roughly spherical shape (**Supplemental Fig. 5B**).

**Fig 5.**
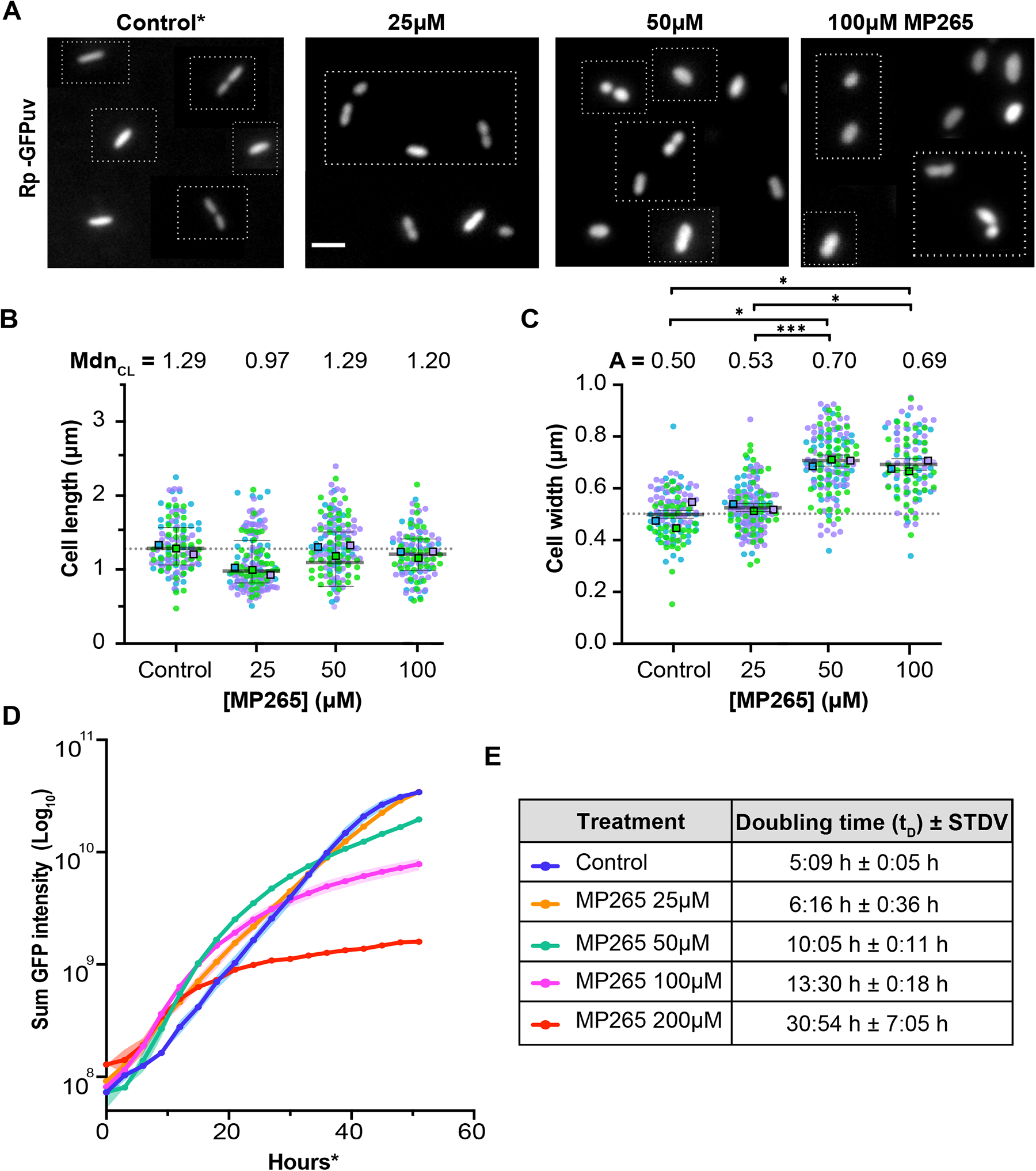
MreB is required to maintain cell width and support robust growth in *R. parkeri.* **(A)** Epifluorescence micrographs of fixed A549 cells infected with Rp-GFPuv and treated with various concentrations (25, 50, 100μM) of MP265 for 24 h. *Control = untreated. Scale bar = 2 µm. **(B)** Cell length and **(C)** cell width distributions of fixed Rp-GFPuv imaged at 48 hours post infection (hpi). Colored dots indicate three independent biological replicates. A Kruskal Wallis with Dunn’s posttest was performed to compare the cell length for all the groups (timepoints) compared to each other, whereas a Welch and Brown-Forsythe ANOVA test with a Dunnett’s posttest was used to compare the cell widths for all the groups (timepoints) compared to each other. *** < P 0.001. **(D)** Semi-log plot of sum GFP intensity over time for A549 cells infected with Rp-GFPuv and treated with the indicated concentration of MP265. **(E)** Doubling times with the indicated treatments. T_D_ indicates the doubling time of Rp-GFPuv or Rp-GFPAa in 24-well plates.

We noticed that cells treated with 50 and 100 µM MP265 not only displayed a significant morphological defect but the number of *R. parkeri* cells per eukaryotic cell was significantly reduced. To understand whether inhibition of MreB affects the growth kinetics of *R. parkeri*, we infected A549 cells with Rp-GFPuv, treated with 0, 25, 50, 100 or 200 µM MP265 and imaged GFP fluorescence every 3 h for ∼55 h. We found that *R. parkeri* cells treated with 25 µM MP265 grew slightly slower than control (untreated) cells (t_D_ control = 5:09 h ± 0:05 h in comparison to t_D_ 25 µM = 6:16 h ± 0:36 h). Strikingly, *R. parkeri* cells treated with 50, 100 or 200 µM MP265 displayed a significant growth defect in comparison to control untreated cells (t_D_ at 50, 100, and 200 µM = 10:05 h ± 0:11 h, 13:30 h ± 0:18 h, and 30:54 h ± 7:05 h, respectively) (**Fig 5DE**) and stopped growing exponentially well before the untreated control. Our newly developed quantitative assays demonstrate that MreB is required for both cell width maintenance and robust growth in *R. parkeri*. Interestingly, in *Rickettsia* species, the elongasome regulator RodZ, which binds MreB and links it to the PG enzymes RodA and PBP2 in *E. coli*, lacks the periplasmic domain that is required to couple MreB and PG dynamics (56). How this truncation of RodZ impacts elongasome dynamics and function in *Rickettsia* species is unclear, but may reflect adaptation of elongasome function to the intracellular environment of a host.

### Conclusions and outlook

In an era of antibiotic resistance, understanding essential biological processes in obligate intracellular human pathogens is of utmost importance, but requires the development of new tools. In our study, we developed and applied quantitative methods to assess peptidoglycan chemistry, cell morphology, cell cycle status, and growth kinetics of the representative SFG pathogen, *R. parkeri*. Using these tools, we established a fundamental understanding of *R. parkeri* morphogenesis in cultured epithelial cells. Moving forward, we can begin to apply these tools to assess contributions of bacterial pathways or host factors into shaping *R. parkeri* cell biology and, by extension, its growth and replication. Though developed in *R. parkeri,* all of the tools described above are translatable to other members of the Rickettsiales, and similar strategies may be undertaken in other obligate intracellular bacteria.

## MATERIALS AND METHODS

### Cell culture

A549 human lung epithelial and Vero monkey kidney epithelial cell lines were kindly provided by Dr. Rebecca Lamason lab (MIT, Cambridge, MA). All cells were grown at 37 °C, 5% CO_2_, and 90% relative humidity. A549 cells were grown in DMEM medium (Gibco, 11965-118) with high glucose (4.5 g/L), containing L-Glutamine and supplemented with 10% heat-inactivated fetal bovine serum (Hi-FBS). Hi-FBS was achieved by placing the FBS (Atlas Biologicals, FP-0500-A) in a water bath at ∼56 °C for 30 minutes. Vero cells were grown in DMEM medium (Gibco, 11965-118) with high glucose (4.5 g/L), containing L-Glutamine and supplemented with 5% Hi-FBS. All cell lines were confirmed to be mycoplasma negative by 4,6-diamidino-2-phenylindole (DAPI) staining. All eukaryotic cell lines were thawed and used after growth rates were established (∼10 passages) and discarded after ∼50 passages.

### Genomic DNA isolation

*R. parkeri* genomic DNA was prepared using the Qiagen DNeasy Blood and Tissue kit following the manufacturer’s protocol. *R. parkeri* (∼1.1 x 10^7^ pfu/mL) were carefully transferred to a 1.5 mL Eppendorf tube. Cells were centrifuged at ∼ 5,000 x *g* and the supernatant was discarded. The bacterial pellet was resuspended in 180 µL ATL Buffer (Qiagen DNeasy Blood and Tissue kit). To lyse the bacterial cells, 20 µL proteinase K were added, mixed thoroughly by vortexing, and incubated at 56 °C for 1 hour. To obtain RNA-free genomic DNA, 4 µL RNAse A (100 mg/mL) were added, mixed by vortexing, and incubated for 2 minutes at room temperature. After the 2-minute incubation, the mixture was vortexed for 15 seconds followed by the addition of 200 µL Buffer AL and 200 µL (96-100%) ethanol. The mixture was immediately vortexed and added into DNeasy Mini spin column. The columns were centrifuged at ∼6,000 x *g* for 1 minute. The flow-through and column were discarded. Next, the DNeasy Mini spin column was placed in a new 2 mL collection tube and 500 µL Buffer AW1 was added. The columns were centrifuged for 1 minute at ∼6,000 x *g.* The flow-through and collection columns were discarded. Next, DNeasy Mini spin column was placed in a new 2 mL collection tube and 500 µL Buffer AW2 was added and centrifuged for 3 minutes at ∼20,000 x *g.* The flow-through and collection tubes were discarded. After discarding the flow-through, the columns were centrigued at ∼20,000 x *g* for 1 minute. Lastly, the DNeasy Mini spin column was transferred into a clean 1.5 mL Eppendorf tube and 100 µL water were directly added into the DNeasy membrane, incubated at room temperature for 1 minute and centrifuged for 1 minute at ∼6,000 x *g*.

### Plasmid and strain construction

#### AausFP1 (GFPAa)

All oligos and plasmids used are listed in Supplemental Table 2,4 and strains are listed in Supplemental Table 3. A plasmid (pRAMEG03) for constitutive expression of cytoplasmic AausFP1 (GFPAa) was constructed as follows: P*_rpsL_*-*arr2* was PCR-amplified from pRAM18dRGA (57) using oligos oEG1544 and oEG1545, digested with PstI and ligated into similarly digested pRAM18dRGA to generate pRAMEG01 (this removes GFPuv from pRAM18dRGA-P*_ompA_*-GFPuv (58) and introduces additional restriction sites). A sequence bearing P*_ompA_* driving *mScarlet* (codon-optimized for *Rickettsia*) followed by a transcriptional terminator was synthesized by IDT. P*_ompA_*-mScarlet-terminator was subcloned into pRAMEG01 using ApaI and StuI to generate pRAMEG02. We attempted to visualize mScarlet in *R. parkeri* cells bearing this plasmid, but signal was too low to be detected. GFPAa codon-optimized for *Rickettsia* was synthesized by IDT and used as a template for PCR using oEG1548 and oEG1549, digested with BsiWI and NgoMIV and ligated into similarly digested pRAMEG02 to generate pRAMEG03.

#### TagBFP

To generate a plasmid for constitutive expression of cytoplasmic TagBFP, oEG1678 and oEG1679 were used to PCR-amplify TagBFP from pRAM18dRA-2xOmpAprCOTBFP (58). The resulting PCR product and pRAMEG02 were digested with BsiWI and NgoMIV and ligated to yield pRAMEG04. To increase the cytoplasmic TagBFP signal, a second copy of TagBFP was introduced into pRAMEG04 by PCR-amplifying P*_ompA_*-TagBFP-terminator using oEG1698 and oEG1699, with pRAMEG04 as the template. The P*_ompA_*-TagBFP-terminator PCR product and pRAMEG04 were digested with SacII and XmaI and ligated to generate pRAMEG05.

#### ZapA-mNeonGreen

To construct a plasmid encoding *zapA-mNeonGreen* (mNG) driven by the P*_zapA_* promoter, *zapA* was PCR-amplified from *R. parkeri* genomic DNA using oEG1568 and oEG1569 and digested with NdeI and NheI. A custom plasmid (pEG1959, pUC-IDT AmpR P*_ompA_*-mNG-ompA3’UTR) was synthesized by IDT for the expression of mNG tagged genes from P*_ompA_* with the *ompA* terminator. mNG was PCR-amplified from this plasmid using oEG1644 and oEG1645 and digested with NheI and NotI. pUC-IDT AmpR P*_ompA_*-mNG-ompA3’UTR was digested with NdeI and NotI and was used as the backbone in a triple-ligation with the *zapA* and mNG fragments. The resulting plasmid (pEG1963) was digested with SacI and NdeI and ligated with P*_zapA_*, which was PCR-amplified from *R. parkeri* genomic DNA using oEG1672 and oEG1673 and digested with the same enzymes. This plasmid was then digested with SacI and SacII to obtain the P*_zapA_*-*zapA-mNG*-terminator fragment, which was then ligated in to similarly digested pRAMEG05 to generate pRAMEG07.

### *R. parkeri* transformation

Bacteria were transformed with pRAMEG03, pRAMEG05, or pRAMEG07 using a small-scale electroporation protocol as described previously (59), except *R. parkeri* infections were scaled up using larger flasks (T175 cm^2^). Briefly, to transform using metabolically active bacteria, T175 cm^2^ flasks of confluent Veros cells were infected for ∼4 days (∼80% of eukaryotic cell rounding) with WT *R. parkeri*. Infected Vero cells were scraped into the growth media, collected in 50 mL conical tubes, and centrifuged at 1700 x g for 5 minutes at 4°C in a JS-7.5 rotor, and resuspended in 3 mL cold K-36 buffer (0.05 M KH_2_PO_4_, 0.05 M K_2_HPO_4_, pH 7, 100 mM KCl and 15 mM NaCl). To release intracellular *R. parkeri*, a “mechanical bead disruption” method was performed as described previously (59) by transferring the homogenate into a 15 mL conical tube containing ∼3 g of 1 mm glass beads. The eukaryotic cells were mechanically lysed by vortexing the 15 mL conical tube on setting 7 using a Pulsing Vortex Mixer (Fisher Scientific Cat # 02215375), with two 30 second pulses and 30 second ice incubations after each pulse. The homogenate was next transferred to (2) 1.5 mL Eppendorf tubes and the host cell debris was pelleted by centrifuging the tubes for 5 minutes at 200 x g. To pellet the bacteria, the supernatant was transferred to (2) 1.5 mL Eppendorf tubes and centrifuged at max speed 16,300 x g for 2 minutes. The pellets were resuspended in 1 mL BHI and frozen at -80 °C. The pellets were resuspended in 1 mL BHI and frozen at -80 or else used for electroporation (below).

To prepare electrocompetent bacteria, these were washed three times with 1 mL cold, filter-sterilized 250 mM sucrose and centrifuged at max speed (16,300 x g) for 2 minutes. Finally, the bacterial pellet was resuspended in 50 μL cold 250 mM sucrose, mixed with 5-10 µg DNA, and electroporated at 1.8 kV, 200 ohms, 25 μF, 5 ms in a pre-chilled 0.1 cm gap cuvette using a MicroPulser (Bio-Rad). Bacteria were recovered in cold 600 µL BHI medium. Next, the media of confluent Vero cells grown in 6-well plates was aspirated, cells were washed with 1 mL phosphate-buffered saline (PBS) and infected with 100 μl of electroporated bacteria (per well). Eukaryotic cells infected with electroporated bacteria were placed in a secondary container and the infection was incubated for 30 minutes at 37°C in a waving shaker. Next, an overlay of DMEM (high glucose, +L-Glutamine) with 2% Hi-FBS and 0.5% (w/v) UltraPure agarose (Invitrogen, 16-500) was added to each well. Plates containing infected Vero cells were incubated at 33°C, 5% CO_2_ in a high humidity chamber. To select for positive transformants, a second agarose overlay containing DMEM (high glucose, +L-Glutamine) with 2% Hi-FBS 0.5% (w/v) UltraPure agarose (Invitrogen, 16-500) and 200 ng/mL rifampicin was added 16 – 24 h after infection. The plates were incubated at 33°C, 5% CO_2_ for 5-7 days in a high humidity chamber.

### *R. parkeri* standard strain expansion, preparation, and enumeration

∼5-7 days after electroporation of *R. parkeri*, ∼6-10 visible plaques were picked, resuspended ∼200µL BHI, and stored at -80°C. To expand the plaques, confluent 6-well plates containing Vero cells were infected at 37°C for 30 min in a waving shaker. Upon infection, each well was supplemented with DMEM (high glucose) 2% Hi-FBS and 200ng/mL rifampicin. Infections were allowed to progress until ∼80% of the eukaryotic cells were infected (∼4-5 days). Bacteria were isolated by a mechanical bead disruption method (as described above). Strains producing fluorescent proteins were confirmed by infecting (in parallel) a second 6-well plate MatTek plate (No. 1.5 Coverslip, 20 mm glass diameter, uncoated plates, part no. P06G-1.5-20-F) containing Vero cells and performing an imaging-based screening. For the imaging-based screening, the infections were allowed to proceed in DMEM 5% FBS supplemented with 200ng/mL rifampicin for ∼24-48 hours. To further expand strains, ∼100µL of bacteria were used to infect either (2) T75 cm^2^ or T175 cm^2^ flasks (for details see below).

To enumerate bacteria, the expanded strains were serially diluted in BHI. Confluent Vero cells grown in 6-well plates were infected with the serially diluted bacteria for 30 minutes at 37 °C using a waving shaker. Infected Vero cell monolayers were immediately overlaid with DMEM 2% Hi-FBS 0.5% (w/v) UltraPure agarose (Invitrogen, 16-500). *R. parkeri* strains harboring the pRAM18dRGA plasmid were overlaid with a second layer of DMEM 2% Hi-FBS 0.5% (w/v) UltraPure agarose containing 200 ng/mL rifampicin ∼16-24 hours after. Plaques were visualized and enumerated 5-7 days post infection to determine the plaque-forming units (PFU) per mL.

### WT *R. parkeri, R. parkeri* GFPuv and *R. parkeri* GFPAusFP1 expansion and enumeration

Parental *R. parkeri* Portsmouth strain (WT *R. parkeri*) and *R. parkeri* expressing GFPuv (pRAM18dRGA-P*_ompA_*-GFPuv) were kindly provided by Dr. Rebecca Lamason (MIT)(58). WT *R. parkeri* and strains producing either GFPuv or GFPAusFP1 (GFPAa) were expanded by infecting 15-20 T175 cm^2^ flasks of confluent Vero cells grown in DMEM (high glucose, +L-Glutamine) medium supplemented with 2% Hi-FBS at a multiplicity of infection (MOI) of about 0.05-0.1. Infected flasks were incubated until ∼80% of the cells were infected, determined by visual inspection (∼4-5 days). *R. parkeri* strains were grown in DMEM (high glucose, +L-Glutamine) and 2% Hi-FBS supplemented with 200 ng/mL rifampicin. To isolate bacteria, infected eukaryotic cells were scraped into the growth media, collected in cold centrifuge bottles, and centrifuged at 12,000 x g for 30 minutes at 4 °C. The cell pellet was resuspended in ∼40 mL cold K-36 buffer (0.05 M KH_2_PO_4_, 0.05 M K_2_HPO_4_, pH 7, 100 mM KCl and 15 mM NaCl) and transferred to a cold dounce 40 mL homogenizer. To release intracellular *R. parkeri*, the host cells were dounce-homogenized for 40-60 strokes. The lysate was subsequently centrifuged at 200 x g for 5 minutes at 4 °C to remove the eukaryotic cell debris. To pellet the bacteria, the supernatant was ultracentrifuged at 58,300 x g for 30 minutes at 4 °C in an SW-27 swinging-bucket rotor. The pellet was resuspended in 1 mL of cold brain heart infusion (BHI) broth (VWR, cat. no. 90003-040) per infected T175 cm^2^ flask, aliquoted, and frozen at −80 °C. Freeze-thaw cycles were minimized to one per stock.

*R. parkeri* strains producing TaqBFP or BFP-ZapA-mNG were expanded following the initial electroporation and imaging-based screening. From the original plaque “mechanical bead disruption” (200 uL), ∼100µL of bacteria were used to infect (2) T75 cm^2^ of confluent Vero cells. Infections were incubated in DMEM (high glucose, +L-Glutamine) medium with 2% Hi-FBS supplemented with 200 ng/mL rifampicin. Infected cells were collected, mechanically disrupted, resuspended in 750µL BHI and stored at – 80C (passage 2, P2). Next, passage 2 was further expanded as described above, except P3 expansion was scaled up using larger flasks (T175 cm^2^). Briefly, (2) confluent T175 cm^2^ flasks of confluent Vero cells were infected with ∼100µL of bacteria and incubated until ∼80% of the cells were infected. Infected cells were collected, mechanically disrupted, resuspended in 1mL BHI and stored at -80C.

### Peptidoglycan analysis

For the muropeptide composition analysis (**Fig 1**), Vero cells were grown in four groups of 20 T175 cm^2^ flask. Two groups were infected with parental WT *R. parkeri* at an MOI of about 0.05-0.1 and incubated at 33 °C with 5% CO_2_ for ∼4 days. The other two uninfected groups were also incubated at 33 °C for 4 days. Cells were harvested using the cell propagation method described above. Briefly, the cells were scraped into the media, collected in centrifuge bottles, and centrifuged at 12,000 x g for 30 minutes at 4 °C. The cell pellet was resuspended in ∼40 mL cold K-36 buffer, transferred to a cold dounce homogenizer, and dounce-homogenized for 40 strokes. The homogenate was centrifuged at 200 x g for 5 min at 4 °C to remove the eukaryotic cell debris. To pellet *R. parkeri*, the supernatant was ultracentrifuged at 58,300 x g for 30 minutes at 4 °C in an SW-27 swinging-bucket rotor. Each of the resulting five cell pellets were resuspended in 3 mL 0.9% NaCl.

UPLC analysis was performed as described previously (31, 32). In brief, pellets resuspended in 3 mL 0.9% NaCl and were boiled in 3 mL SDS 5% for 2 h. Sacculi were repeatedly washed with MilliQ water by ultracentrifugation at ∼540,000 x g for 15 min at 20 °C in a TLA-100.3 rotor (OptimaTM Max ultracentrifuge, Beckman) until completely removing SDS traces. The samples were then treated with muramidase (100 µg/mL) for 16 hours at 37 °C. Muramidase digestion was stopped by boiling and coagulated proteins were removed by centrifugation at 14,000 rpm for 10 minutes in a tabletop centrifuge. The supernatants were first adjusted to pH 8.5-9.0 with sodium borate buffer and then sodium borohydride was added to a final concentration of 10 mg/mL. After reduction during 30 minutes at room temperature, the pH of the samples was adjusted to pH 3.5 with orthophosphoric acid. UPLC analysis of muropeptides was performed on a Waters UPLC system (Waters Corporation, USA) equipped with an ACQUITY UPLC BEH C18 Column, 130Å, 1.7 µm, 2.1mm X 150mm (Waters, USA) and a dual wavelength absorbance detector. Elution of muropeptides was detected at 204 nm. Muropeptides were separated at 45 °C using a linear gradient from buffer A (formic acid 0.1% in water) to buffer B (formic acid 0.1% in acetonitrile) in an 18-minute run, with a 0.25 mL/min flow. Muropeptide identity was confirmed by MS/MS analysis, using a Xevo G2-XS QTof system (Waters Corporation, USA).

After the identification of all the relevant muropeptides by MS/MS, we calculated the relative molar abundances of the major muropeptides of two biological replicates (20 T175 cm^2^ per biological replicate).

### Epifluorescence and phase microscopy and image analysis

A549 (**Fig 2**) and Vero (**Supplemental Fig 1**) cells were grown in 35 mm MatTek dishes (No. 1.5 uncoated coverslip, 20 mm glass, part no. P35G-1.5-20-C.S) for 1-2 days (until reaching 80% confluency) and infected with *R. parkeri* producing GFPuv at an MOI of 0.25-0.5. The infection was incubated at 33°C with 5% CO_2_ for the duration of the experiment. The dishes were imaged with a Nikon Eclipse Ti inverted microscope through a Nikon Plan Fluor 100 × (numeric aperture, 1.30) oil Ph3 objective and GFP filters with a Photometrics CoolSNAP HQ2 cooled charge-coupled-device (CCD) camera and Nikon Elements Imaging Software. Images of three biological replicates were collected at 6, 12, 24, and 48 hours post-infection (hpi) in stage incubation chamber (Tokai Hit) and an STX Stage Top Incubator (Tokai Hit), where the top lid temperature was set at 56 °C, chamber temperature at 33 °C, and 5% CO_2_,

Quantitative evaluation of the morphology of live or fixed *R. parkeri* (**Supplemental Fig. 1**, **Fig. 5B**) was performed in the FIJI (60) plug in, MicrobeJ (61), by using the fluorescence signal of raw images to define the outline of the cells. To ensure an efficient analysis in MicrobeJ, unfocused bacterial cells were removed from the Z-stack slices prior to starting the analysis. To determine the cell length and width distributions, we measured the fluorescent cell profiles using MicrobeJ. Superplots were constructed in Adobe Illustrator by superimposing the three scatter plot replicates with their respective median length or average width (43). To get the global median cell length or average width, the cells from the three replicates were combined and a fourth scatter plot was constructed and superimposed. The demograph in **Fig 3B** was created by plotting the cell profiles and sorting the cells by their length. To quantify the percentage of constricting cells (**Fig 3C**), raw images of cells producing GFPuv were quantified by counting the number of pre-divisional cells over the total number of cells. Three independent biological replicates were quantified.

Lastly, for **Fig 3D** A549 cells were grown 24-well MatTek plates (No. 1.5 uncoated coverslip, 13 mm glass) 1-2 days (until reaching 80% confluency) and infected with *R. parkeri* producing cytoplasmic BFP and ZapA-mNG. The plate was imaged with a Nikon Eclipse Ti inverted microscope through a Nikon Plan Fluor 100 × (numeric aperture, 1.30) oil Ph3 objective, GFP and BFP filters with a Photometrics CoolSNAP HQ2 cooled charge-coupled-device (CCD) camera and Nikon Elements Imaging Software. Images of three biological replicates were collected at 24hours post-infection (hpi) in stage incubation chamber (Tokai Hit) and an STX Stage Top Incubator (Tokai Hit), where the top lid temperature was set at 56 °C, chamber temperature at 33 °C, and 5% CO_2_.

### Quantitative validation of the morphology of *R. parkeri* using phase contrast microscopy

Quantitative validation of *R. parkeri* morphology measurements (**Supplemental** **Fig. 2**) were performed by first growing (3) T175 cm^2^ flasks of Vero cells in DMEM (high glucose, +L-Glutamine) medium supplemented with 5% Hi-FBS. Vero cells were subsequently infected with WT *R. parkeri* (parental) or parental *R. parkeri* producing GFPuv (Rp-GFPuv) at multiplicity of infection (MOI) of 0.05-0.1. The infection was incubated at 33 °C with 5% CO_2_ for 48 hours. Bacteria were mechanically lysed by performing a “mechanical bead disruption” method (as described above). After lysis, bacteria were fixed in 70% ethanol for 15 minutes, centrifuged at max speed (16,300 x g) for 2 minutes and resuspended in K-36 buffer. Fixed bacteria were immobilized on 1.25% (w/v) UltraPure agarose pads made with K-36 buffer, and imaged via phase contrast microscopy using a Nikon Eclipse Ti inverted microscope through a Nikon Plan Fluor 100 × (numeric aperture, 1.30) oil Ph3 objective with a Photometrics CoolSNAP HQ2 cooled charge-coupled-device (CCD) camera and the Nikon Elements Software. To quantify cell length and width, phase-contrast images were analyzed using the MicrobeJ plugin for Fiji (61).

### Fluorescent growth curves

To evaluate growth kinetics of *R. parkeri*, A549 (**Fig. 4**) and Vero (**Supplemental Fig. 3**) cells were grown in 24-well MatTek dishes (No. 1.5 uncoated coverslip, 13 mm glass, cat no.) for 1-2 days (or until reaching 95% confluency) and infected with *R. parkeri* producing GFPuv, AausFP1, or TagBFP at an MOI of 0.5-1.2 for 24 hours before imaging. To run the growth curves, a multi-mode imaging plate reader method was developed using the Cytation1 imaging reader (Agilent, Biotek). Phase-contrast and fluorescence images were captured through the Olympus Plan Fluorite 20× objective (numeric aperture, 0.45) with a 465 nm LED GFP or 365 nm LED BFP filter cubes, and focused using a laser autofocus cube (Agilent BioTek 1225010). Images were collected every 3 hours for ∼55 h and the infection was incubated at 33 °C and 5% CO_2_. Using the Gen5 software, an automated protocol was developed in which 9 images and 10 slices were obtained at the center of wells. These images were subsequently crop stitched, linearly blended, transformed by background flattening, and Z-projected by using a focus stacking method. Next, a cellular analysis was conducted on the Z-projected images to allow for calculation of the sum of GFP or BFP intensity.

### PFU growth curves

To evaluate growth kinetics of *R. parkeri* producing GFPuv or GFPAa in Vero cells by plaque assays (**Supplemental Fig. 3C**), cells were grown in T75 cm^2^ flasks and infected with Rp-GFPuv or Rp-GFPAa. To quantify the number of infectious bacteria, infected Vero cells were harvested from the T75 cm^2^ flasks, and serial dilutions were performed in cold BHI. Vero cells grown in 6-well plates were then infected and the infection was incubated for 30 minutes at 37°C. DMEM 2% Hi-FBS 0.5% Ultrapure agarose (Invitrogen) was used to overlay the wells. A second layer of DMEM 2% Hi-FBS 0.5% Ultrapure agarose containing 200 µg/mL rifampicin was added ∼16 h after. The plaques were quantified 5-7 days post infection.

### Global complex cell shape analysis

For the cell shape analysis on **Fig 5B** and **Supplemental Fig 5**, binary masks of *R. parkeri* producing cytoplasmic GFP were made and loaded into the Celltool software (55). Cell shapes or “polygonal contours” of control cells as well as cells treated with 0, 25, 50 or 100 µM MP265 were extracted from the binaries. To take into consideration the shape variability in the MP265-perturbed cells, a model for all the contours was made. 95% of the shape variance was accounted to “shape mode 1”, which roughly reflects cell length and “shape mode 2”, which roughly reflects cell width. To understand the impact of increasing concentrations of MP265 on *R. parkeri* cell shape, a two-dimensional principal component analysis (PCA) of shape and box-plot distributions were constructed. Box-plots were constructed in Prism 9.0.

### Statistical analysis, experimental variability, and reproducibility

One-way analysis of variance (ANOVA) Kruskal-Wallis test with Dunn’s posttest was used to the cell lengths, whereas a Welch and Brown-Forsythe ANOVA test with a Dunnets posttest was used to compare the cell widths for all the groups (timepoints) to each other. All statistical analyses were performed in Prism 9.0.

## ACKNOWLEDGEMENTS

We thank the members of the Goley lab for helpful discussions and input. We thank Dr. Rebecca Lamason, Dr. Matthew Welch, and the Lamason and Welch laboratories for strains, protocols, training, and helpful discussions about the *R. parkeri* biology. We thank Dr. Nathan Shaner for helpful discussions regarding fluorescent proteins. We also thank Dr. Allison Cross from Agilent for her plate reader guidance and protocol optimization. This work is funded in part by the NIH through awards R21AI159075 (to E.D.G.) and T32GM144272 (training grant support of E.S. and M.F.), and by the National Science Foundation Postdoctoral Research Fellowship in Biology through PRFB award number 2010706 (W.M.F.C). Research in the Cava lab is supported by the Swedish Research Council, the Laboratory for Molecular Infection Medicine Sweden (MIMS), Umeå University, the Knut and Alice Wallenberg Foundation (KAW) and the Kempe Foundation.

