## Supplemental Figures and Legends for "Quantitative analysis of morphogenesis and growth dynamics in an obligate intracellular bacterium"

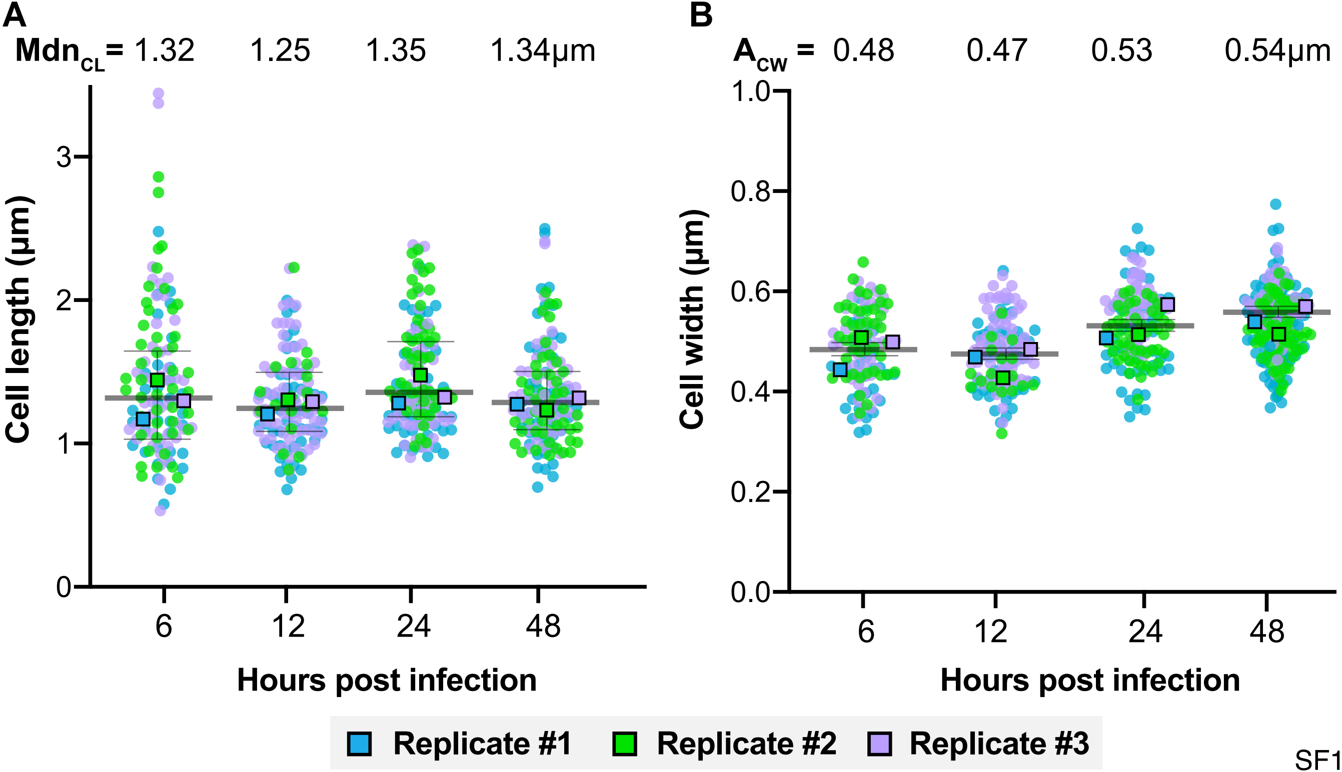
A
$Mdn_{CL}$ = 1.32 1.25 1.35 1.34µm
Cell length (µm)
0
1
2
3
6
12
24
48
Hours post infection
B
$A_{CW}$ = 0.48 0.47 0.53 0.54µm
Cell width (µm)
0.0
0.2
0.4
0.6
0.8
1.0
6
12
24
48
Hours post infection
Replicate #1
Replicate #2
Replicate #3

SF1

**Supplemental Fig 1. Quantitative evaluation of the morphology of *R. parkeri* in green monkey epithelial (Vero) cells. (A)** Cell length or (B) distributions of live Rp-GFPuv imaged at indicated hours post infection (hpi). Colored dots (blue, purple, green) indicate three independent biological replicates, whereas colored boxes represent independent median cell length ($Mdn_{CL}$) or average cell width ($A_{CW}$). A Kruskal Wallis with Dunn's post-test was performed to compare the cell length for all the groups (time points) compared to each other, whereas a Welch and Brown-Forsythe ANOVA test with a Dunnett's post-test was used to compare the cell widths for all the groups (time points) to each other. No statistical significance was found comparing cell length or cell width across time points.

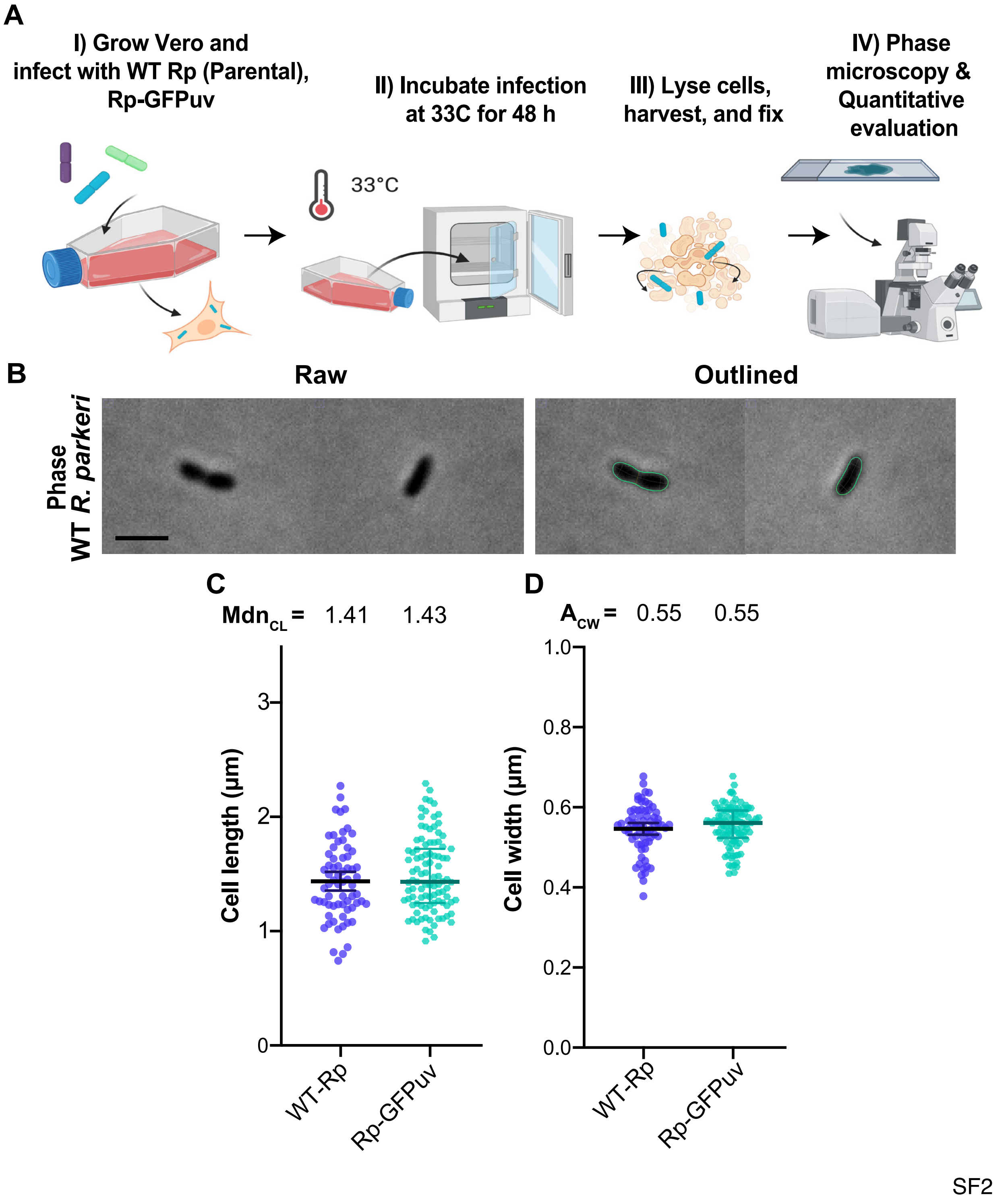
A
I) Grow Vero and infect with WT Rp (Parental), Rp-GFPuv
II) Incubate infection at 33C for 48 h
III) Lyse cells, harvest, and fix
IV) Phase microscopy & Quantitative evaluation
33°C
B
Raw
Outlined
WT R. parkeri Phase
C
Mdn$_{CL}$ = 1.41 1.43
Cell length (µm)
0
1
2
3
WT-Rp
Rp-GFPuv
D
A$_{cw}$ = 0.55 0.55
Cell width (µm)
0.0
0.2
0.4
0.6
0.8
1.0
WT-Rp
Rp-GFPuv

**Supplemental Fig 2. Validation of quantitative evaluation of morphology by phase contrast microscopy. (A)** Strategy for the validation of the *R. parkeri* morphology. **(B)** Raw Rp-GFPuv images obtained via phase contrast microscopy (48 hours post infection, 48 hpi). **(B)** Composite of raw phase contrast images (48 hpi) were used to detect cell outlines ("Outlined") using MicrobeJ software. Representative cells were selected from different images to make a composite. Scale bar = 2 μm. **(C)** Cell length or **(D)** cell width distributions of fixed WT and Rp-GFPuv. N ~100 cells per strain. A Kruskal Wallis with Dunn's post-test was performed to compare the cell length for all the groups (time points) compared to each other, whereas a Welch and Brown-Forsythe ANOVA test with a Dunnett's post-test was used to compare the cell widths for all the groups (time points) to each other. No statistical significance was found comparing cell length or cell width across time points.

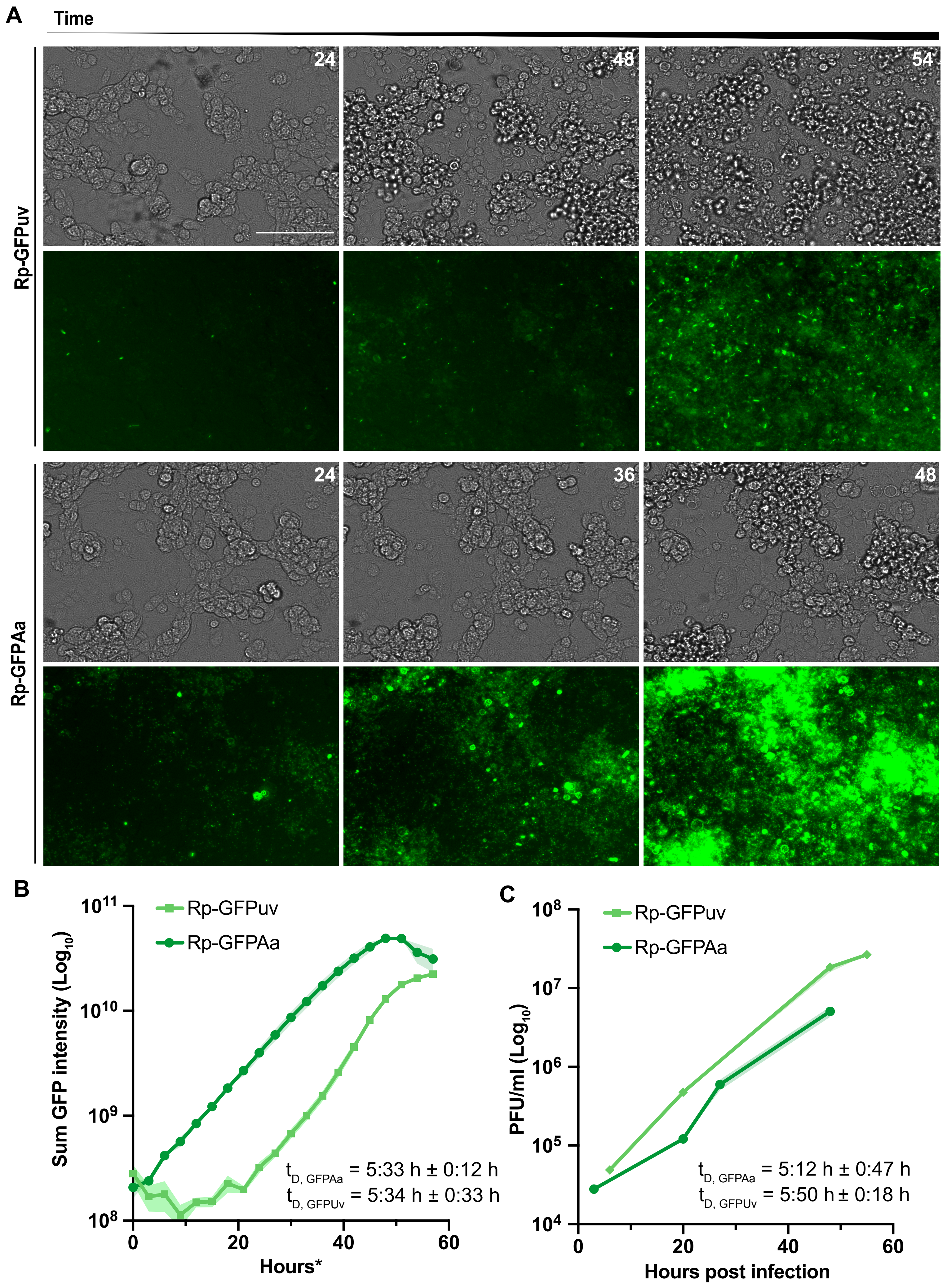
A
Time
24
48
54
Rp-GFPuv
24
36
48
Rp-GFPAa
B
Rp-GFPuv
Rp-GFPAa
Sum GFP intensity (Log10)
10^11
10^10
10^9
10^8
t_D, GFPAa = 5:33 h ± 0:12 h
t_D, GFPUv = 5:34 h ± 0:33 h
0
20
40
60
Hours*
C
Rp-GFPuv
Rp-GFPAa
PFU/ml (Log10)
10^8
10^7
10^6
10^5
10^4
t_D, GFPAa = 5:12 h ± 0:47 h
t_D, GFPUv = 5:50 h ± 0:18 h
0
20
40
60
Hours post infection

SF3

**Supplemental Fig 3. Fluorescence-based quantification of intracellular growth dynamics of *R. parkeri* in Vero cells. (A)** Phase-contrast (top) and GFP (bottom) images taken with the BioTek Cytation1 at indicated hours post imaging. Scale bar = 100µm. **(B)** Semi-log plot of sum GFP intensity over time for Vero cells infected with Rp-GFPuv or Rp-GFPAa. $t_D$ indicates the doubling time of Rp-GFPuv or Rp-GFPAa in 24-well plates. **(C)** Semi-log plot of PFU/mL of the indicated strains over 60 hours of infection. $t_D$ indicates the doubling time of Rp-GFPuv or Rp-GFPAa.

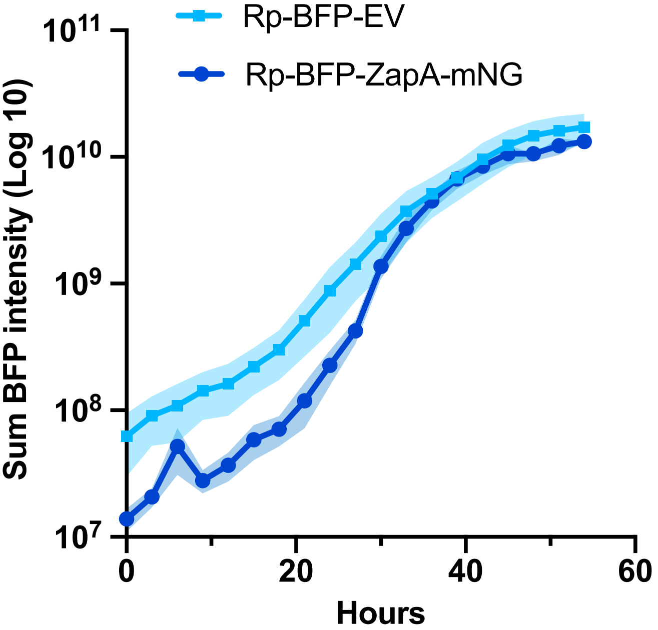


Rp-BFP $t_D$ = 5:55 h ± 0:48 h
Rp-ZapA-mNG $t_D$ = 5.00 h ± 0:27 h

SF4

**Supplemental Fig 4. Fluorescence-based quantification of growth kinetics in *R. parkeri* producing ZapA-mNG.** Semi-log plot of BFP intensity over time for A549 cells infected with *R. parkeri* producing BFP and *R. parkeri* producing both, BFP and ZapA-mNG from the same plasmid. $t_D$ indicates the doubling time of Rp-BFP or Rp-BFP, ZapA-mNG.

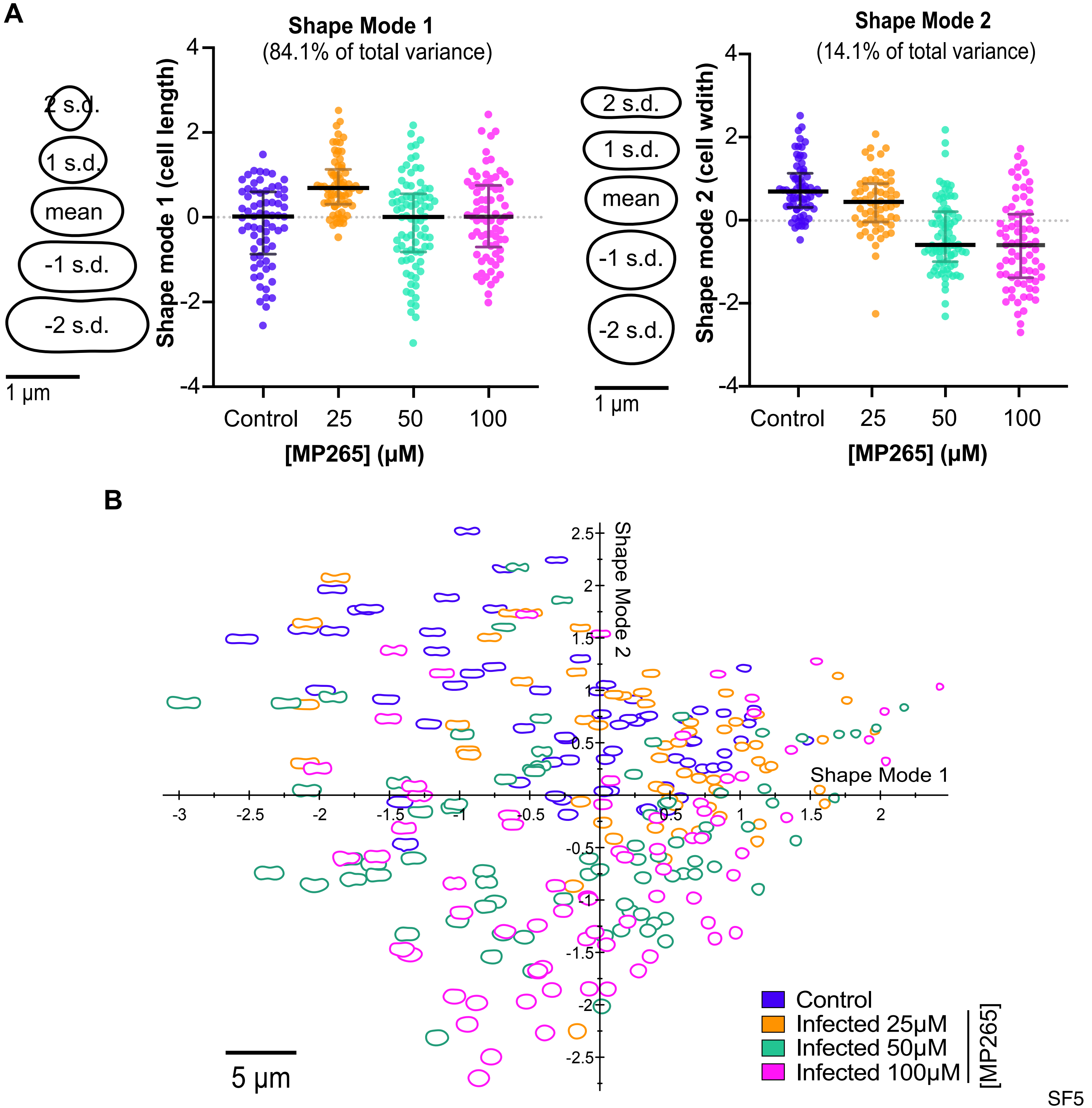
A
Shape Mode 1
(84.1% of total variance)
2 s.d.
1 s.d.
mean
-1 s.d.
-2 s.d.
1 µm
Shape mode 1 (cell length)
Control
25
50
100
[MP265] (µM)
Shape Mode 2
(14.1% of total variance)
2 s.d.
1 s.d.
mean
-1 s.d.
-2 s.d.
1 µm
Shape mode 2 (cell wdith)
Control
25
50
100
[MP265] (µM)
B
Shape Mode 2
Shape Mode 1
5 µm
Control
Infected 25µM
Infected 50µM
Infected 100µM
[MP265]

SF5

**Supplemental Fig 5. Principal component analysis of *R. parkeri* treated with MP265. (A)** Scatter plots of shape mode 1 (approximately reflecting cell length) and shape mode 2 (approximately representing cell width) values for Rp-GFPuv cells in A549 host cells untreated (control) or treated with the indicated concentration of MP265 for 24 h (N = 100). **(B)** Shape mode 1 plotted against shape mode 2 from principal component analysis (PCA) of Rp-GFPuv treated with various concentrations of MP265 after 24 hours. A Kruskal Wallis with Dunn's posttest was performed to compare the medians of shape mode, whereas a Welch and Brown-Forsythe ANOVA test with a Dunnett's posttest was used to compare the means of shape mode 2 for all the groups. *** < P 0.001.

**Supplemental Video 1. Fluorescence-based quantification of intracellular growth dynamics of *R. parkeri* in A549 cells.** Phase-contrast time-lapse of A549 cells infected with *R. parkeri* producing GFPuv over time.

**Supplemental Video 2. Fluorescence-based quantification of intracellular growth dynamics of *R. parkeri* in A549 cells.** GFP time-lapse of A549 cells infected with *R. parkeri* producing GFPuv over time.

**Supplemental Video 3. Fluorescence-based quantification of intracellular growth dynamics of *R. parkeri* in A549 cells.** Phase-contrast time-lapse of A549 cells infected with *R. parkeri* producing GFPAusFP1 (GFPAa) over time.

**Supplemental Video 4. Fluorescence-based quantification of intracellular growth dynamics of *R. parkeri* in A549 cells.** GFP time-lapse of A549 cells infected with *R. parkeri* producing GFPAusFP1 (GFPAa) over time.

**Supplemental Video 5. Fluorescence-based quantification of intracellular growth dynamics of *R. parkeri* in A549 cells.** Phase contrast time-lapse of A549 cells infected with *R. parkeri* producing BFP over time.

**Supplemental Video 6. Fluorescence-based quantification of intracellular growth dynamics of *R. parkeri* in A549 cells.** BFP time-lapse of A549 cells infected with *R. parkeri* producing BFP over time.

**Supplemental Video 7. Fluorescence-based quantification of intracellular growth dynamics of *R. parkeri* in A549 cells.** Phase contrast time-lapse of A549 cells infected with *R. parkeri* producing both BFP and ZapA-GFPmNG.

**Supplemental Video 7. Fluorescence-based quantification of intracellular growth dynamics of *R. parkeri* in A549 cells.** BFP time-lapse of A549 cells infected with *R. parkeri* producing both BFP and ZapA-GFPmNG.
